# Identification of interpretable clusters and associated signatures in breast cancer single cell data: a topic modeling approach

**DOI:** 10.1101/2022.09.13.507779

**Authors:** Malagoli Gabriele, Valle Filippo, Barillot Emmanuel, Caselle Michele, Martignetti Loredana

## Abstract

Topic modeling is a popular technique in machine learning and natural language processing, where a corpus of text documents is classified into themes or topics using word frequency analysis. This approach has proven successful in various biological data analysis applications, such as predicting cancer subtypes with high accuracy and identifying genes, enhancers, and stable cell types simultaneously from sparse single cell epigenomics data. The advantage of using a topic model is that it not only serves as a clustering algorithm, but it can also explain clustering results by providing word probability distributions over topics.

Our study proposes a novel topic modeling approach for clustering single cells and detecting topics (gene signatures) in single-cell datasets that measure multiple omics simultaneously. We applied this approach to examine the transcriptional heterogeneity of luminal and triple negative breast cancer cells using patient-derived xenograft models with acquired resistance to chemotherapy and targeted therapy. Through this approach, we identified protein-coding genes and long noncoding RNAs (lncRNAs) that group thousands of cells into biologically similar clusters, accurately distinguishing drug-sensitive and resistant breast cancer types. In comparison to standard state-of-the-art clustering analyses, our approach offers optimal partitioning of genes into topics and cells into clusters simultaneously, producing easily interpretable clustering outcomes. Additionally, we demonstrate that an integrative clustering approach, which combines the information from mRNAs and lncRNAs treated as disjoint omics layers, enhances the accuracy of cell classification.

## INTRODUCTION

The recent development of single cell technologies has enabled us to achieve a highly detailed view of the transcriptome in many biological studies. A fundamental step in the analysis of single cell transcriptome data is the identification of cell populations, i.e. groups of cells that show “similar” expression patterns.

To solve this task, many popular algorithms for unsupervised clustering have been implemented, which still present some computational challenges, such as estimating the optimal number of cell types [1] or biological interpretation and annotation of the identified clusters [2].

A novel approach, based on topic modeling, has been recently introduced for simultaneously discovering clusters and interpretable gene signatures from bulk [3][4] and single cell [5] transcriptomic datasets. Importantly, the algorithm used here for topic modeling, called hierarchical Stochastic Block Modeling (hSBM) [6][7] is able to determine the optimal number of clusters (cell subpopulations) at different levels of resolution (i.e., the algorithm is able to reconstruct the hierarchical organization of cells into clusters) without a priori setting the number of expected clusters and provides the topics (i.e. sets of genes) significantly associated with the identified clusters.

Here, we apply the hSBM algorithm to study the transcriptional heterogeneity of breast cancers and the role of long non-coding RNAs (lncRNAs) in contributing to this heterogeneity. LncRNAs are a large class of transcripts that are expressed in human cells without coding potential [8][9]. In healthy tissues, they have been shown to have lower expression, increased tissue-specificity, and greater expression variability across individuals than protein-coding genes [10][11]. In cancer, lncRNAs are commonly dysregulated and their aberrant expression has been shown to be tumor-type specific [12][13]. However, the expression of lncRNAs at single cell resolution has not yet been extensively studied. It is still unclear whether lncRNAs are highly expressed in subsets of cells within tissues, despite appearing lowly expressed in bulk populations. A single molecule, single cell RNA FISH survey of 61 lncRNAs in three human cell types showed that the low abundance of lncRNAs in bulk population measurements is not due to a small subpopulation of cells expressing lncRNAs at high levels, and overall lncRNA are no different than mRNA in their levels of cell-to-cell heterogeneity [14]. On the other hand, a larger scale study based on Smart-seq-total technology [15] has assayed a broad spectrum of mRNAs and lncRNAs from single cells and showed that the lncRNA content of cells significantly differs across cell types and dynamically changes throughout cellular processes such as cell cycle and cell differentiation. Another large-scale study, combining bulk tissue RNA-seq and scRNA-seq to deeply profile lncRNA expression during neocortical development, found that many lncRNAs are specific to distinct cell types and abundantly expressed in individual cells [16], thus supporting that cell type-specific expression of lncRNAs contributes to the low levels of lncRNAs observed in tissues. Our work corroborates the latter kind of result.

Existing computational methods for analysing single cell RNA-seq (scRNA-seq) data do not consider the differences between protein-coding mRNAs and lncRNAs in terms of data dispersion and sparsity. Unsupervised clustering typically suffers from large fractions of observed zeros. In comparison to the analysis of mRNAs, the computational analysis of lncRNAs at single cell is expected to be more challenging due to their high expression variability [2, 18].

In our study, we used hSBM to investigate the transcriptional heterogeneity of luminal and triple negative breast cancer cells using patient-derived xenograft models of acquired resistance to chemotherapy and targeted therapy [18]. First, we clustered the cells based on the expression of mRNAs and lncRNAs separately. We then used a multi-omics version of hSBM [4] to perform an integrative clustering that combines the information coming from the two RNA families treated as disjoint omics layers. We show that this strategy improves cell classification compared to clustering obtained by investigating mRNAs and lncRNAs separately but also compared to clustering obtained on the expression measures of mRNAs and lncRNAs concatenated in a single matrix.

The main innovation we introduced is a reproducible and unsupervised procedure to investigate the gene content associated with cell clusters. For this purpose, we have developed a method to assign the most specific set of topics to each cluster and query them through gene set functional annotation databases (see Methods). The topics have been interpreted in terms of functionally related gene sets of mRNAs and lncRNAs. This workflow allows for a deep investigation of the biological content of each cluster. The results show a clear enrichment of topics for pathways involved in subtyping and progression of breast cancer and for sets of lncRNA that are interactors of important transcription factors involved in cancer. We have identified some lncRNAs associated with breast cancer cell subpopulations already well known in the literature, such as MALAT1 and NEAT1, and highlighted some others that may be clinically relevant.

## RESULTS

### 1. Clustering of drug-sensitive and resistant breast cancer cells using hSBM

To explore intra-tumor transcriptional heterogeneity in breast cancer, we used a dataset of scRNAseq profiles of patient-derived xenograft (PDX) models of luminal and triple negative breast cancer with acquired drug resistance [18]. After scRNA-seq data pre-processing (see Methods), we obtained two expression matrices containing respectively 1959 mRNAs and 1136 lncRNAs profiled in 3723 cells from drug-sensitive and resistant tumors. We translated each expression matrix into a weighted bipartite network where nodes represent genes (mRNAs or lncRNAs) and cells. Genes are linked to the cells with weighted edges based on their level of expression. We applied the hSBM algorithm to calculate the best partitions of features and cells. The output of hSBM is a hierarchical and probabilistic organization of the network in *blocks* of connected genes and cells. For each level of hierarchy, we obtain three matrices of probability membership, corresponding to the probability of cells to be associated to a cluster (which is a delta function since we set the model to output “hard” clusters), the probability of association of genes to topics, and the probability of association of a topic to a cell. In our context, the output of hSBM informs us about the best partitioning of cells into clusters with similar transcriptional profiles and about which genes are most strongly associated with these clusters.

The partition of the network based on mRNA expression shows three hierarchical levels of cell classification (Figure 1A, B, C). In the hierarchical level 2 (Figure 1C), the cells are classified into two clusters containing respectively the luminal and triple negative tumor cells, regardless of their resistant or drug-sensitive phenotype. The level 1 of clustering resolution (Figure 1B) separates cells into nine clusters, three of which contain a mixture of sensitive and resistant cells, suggesting that heterogeneous populations of sensitive and resistant cells emerged. The largest of these mixed clusters includes 999 triple negative sensitive and resistant cells, while none of the mixed clusters contain both basal and luminal cells. The level 0 of classification (Figure 1A) separates cells into 49 clusters, all containing cells from specific tumor types.

**Figure 1.**
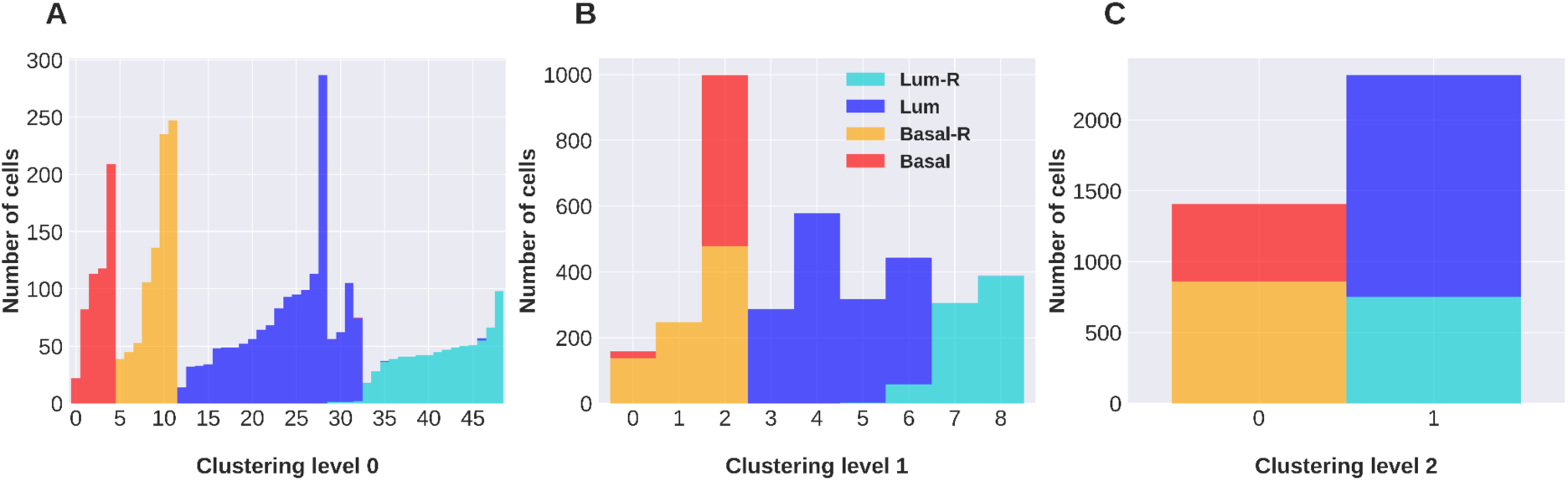
Clustering of cells obtained with hSBM applied to the mRNAs expression dataset. For each hierarchical level of clustering (A, B, C), columns report the size of the identified clusters and the tumor of origin of their component cells.

Also the partition of the lncRNA-based network exhibits three hierarchical levels of cell classification (Figure 2A, B, C). Similar to the clustering of the mRNA network, the level 2 (Figure 2C) classifies cells in two clusters with luminal and basal tumor cells. However, the level 1 and level 0 hierarchical levels show some differences compared to the classification based on mRNAs. The level 1 (Figure 2B) separates cells into seven clusters, all containing a mixture of resistant and sensitive cells. The level 0 of classification (Figure 2A) separates cells into 249 clusters, many of which include both resistant and sensitive cells.

**Figure 2.**
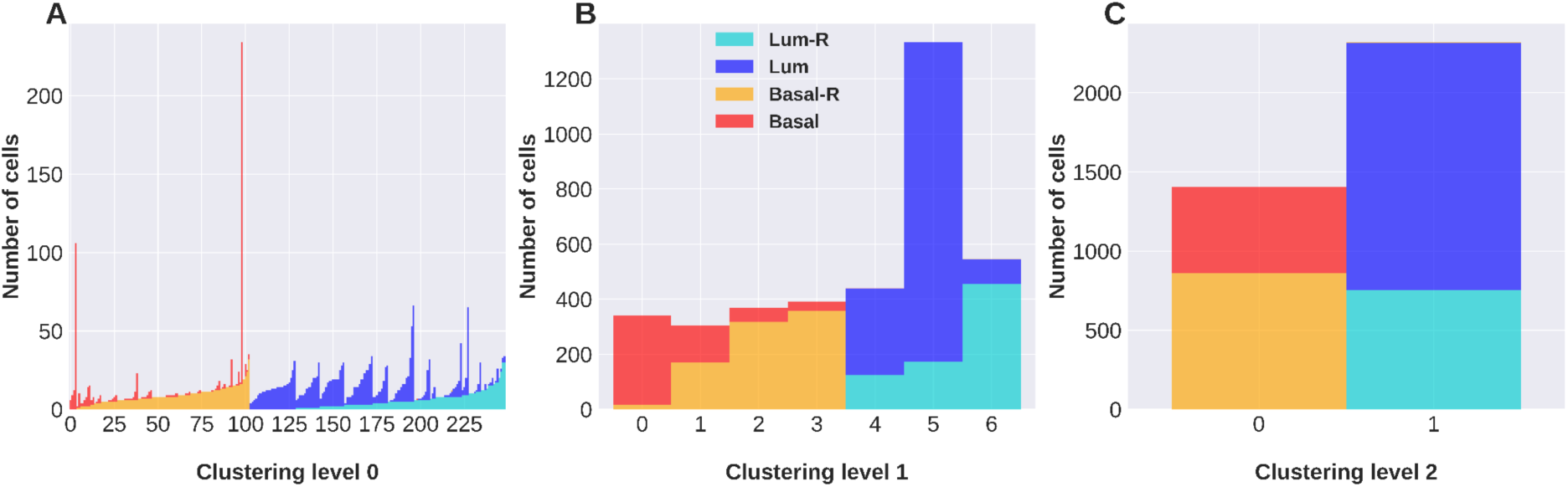
Clustering of cells obtained with hSBM applied to the lncRNAs expression dataset. For each hierarchical level of clustering (A, B, C), columns report the size of the identified clusters and the tumor of origin of their component cells.

To evaluate the performance of the hSBM clustering algorithm, we calculated a score based on the normalized mutual information (NMI) between the partition found by the algorithm and the tumor of origin of the cells (see Methods). The NMI/NMI* scores obtained for the mRNA-based and lncRNA-based clustering are reported in Table 1. The NMI/NMI* score decreases as clustering resolution increases from level 2 to level 0, as cells from the same tumor type are separated into different clusters. At the level 2 and level 1 resolution, NMI/NMI* scores are comparable for clustering obtained with mRNAs and that obtained with lncRNAs, whereas at the level 0 NMI/NMI* is an order of magnitude lower for clustering obtained with lncRNAs. The low NMI/NMI* obtained for the level 0 of clustering with lncRNAs shows that their expression is not very informative of the tumor type at this clustering hierarchical level. Instead, the classification of cells obtained at the level 1 provides information strongly correlated with tumor types. Remarkably, the mRNA-based clustering divides cells differently than lncRNA-based clustering, suggesting that the expression of these two RNA families carries complementary information about cell heterogeneity.

**Table 1.** NMI/NMI* scores computed to assess the consistency between the cell clusters obtained from diverse experiments conducted with hSBM and Seurat algorithms, and the original tumor from which the cells were derived.

| NMI/NMI* | Clustering level 0 | Clustering level 1 | Clustering level 2 |
| --- | --- | --- | --- |
| hSBM-mRNAs | 66 | 325 | 1560 |
| hSBM-lncRNAs | 9 | 359 | 1642 |
| hSBM-mRNAs-lncRNAs | 53 | 294 | 1618 |
| multibranch SBM<br>mRNAs-lncRNAs | 56 | 398 | 1603 |
| Seurat-mRNAs | 465 |  |  |
| Seurat-lncRNAs | 614 |  |  |
| Seurat-WNN<br>mRNAs-lncRNAs | 451 |  |  |

### 2. Cells are better classified by analyzing the expression of mRNAs and lncRNAs as separate omics layers

We then simultaneously use mRNAs and lncRNAs expression data to perform integrative clustering. These data are generated on the same platform in a single experiment, and the standard approach for integrative analysis is to concatenate the expression measurements into a single matrix on which to apply the clustering algorithm. A more sophisticated approach would consider the inherent difference between mRNAs and lncRNAs expression and treat the two RNA families as disjoint omics layers. Indeed, we have verified that in this dataset lncRNAs show statistically higher coefficients of variance than mRNAs (Supp. Figure 1).

To integrate the information coming from mRNAs and lncRNAs, we designed two different experiments. In the first one, we concatenate the mRNAs and lncRNAs expression measurements into a single matrix of 3095 features and apply hSBM algorithm on it. In the second experiment, the mRNAs and lncRNAs matrices are treated as disjoint omics layers and a multi-omics version of hSBM is applied, that is called multibranch SBM [4] and is similar to the multilayer SBM algorithm used for text document analysis [6]. This extends the network-based topic modeling algorithm to a weighted tri-partite network where nodes represent mRNAs, lncRNAs and cells. Similarly to hSBM, mRNAs and lncRNAs are linked to the cells according to their expression level, while no mRNA-lncRNA edges are present, and the best partition of the tripartite network is computed.

The two experiments produce very different results in terms of cell partition (Figure 3). In the first one, based on hSBM applied to the concatenated expression matrices, we obtain eleven clusters almost exclusively composed of cells specific to one specific tumor type (Figure 3A). The multibranch SBM algorithm separates the cells into seven clusters, three of which contain a mixture of resistant and sensitive cells (Figure 3B).

**Figure 3.**
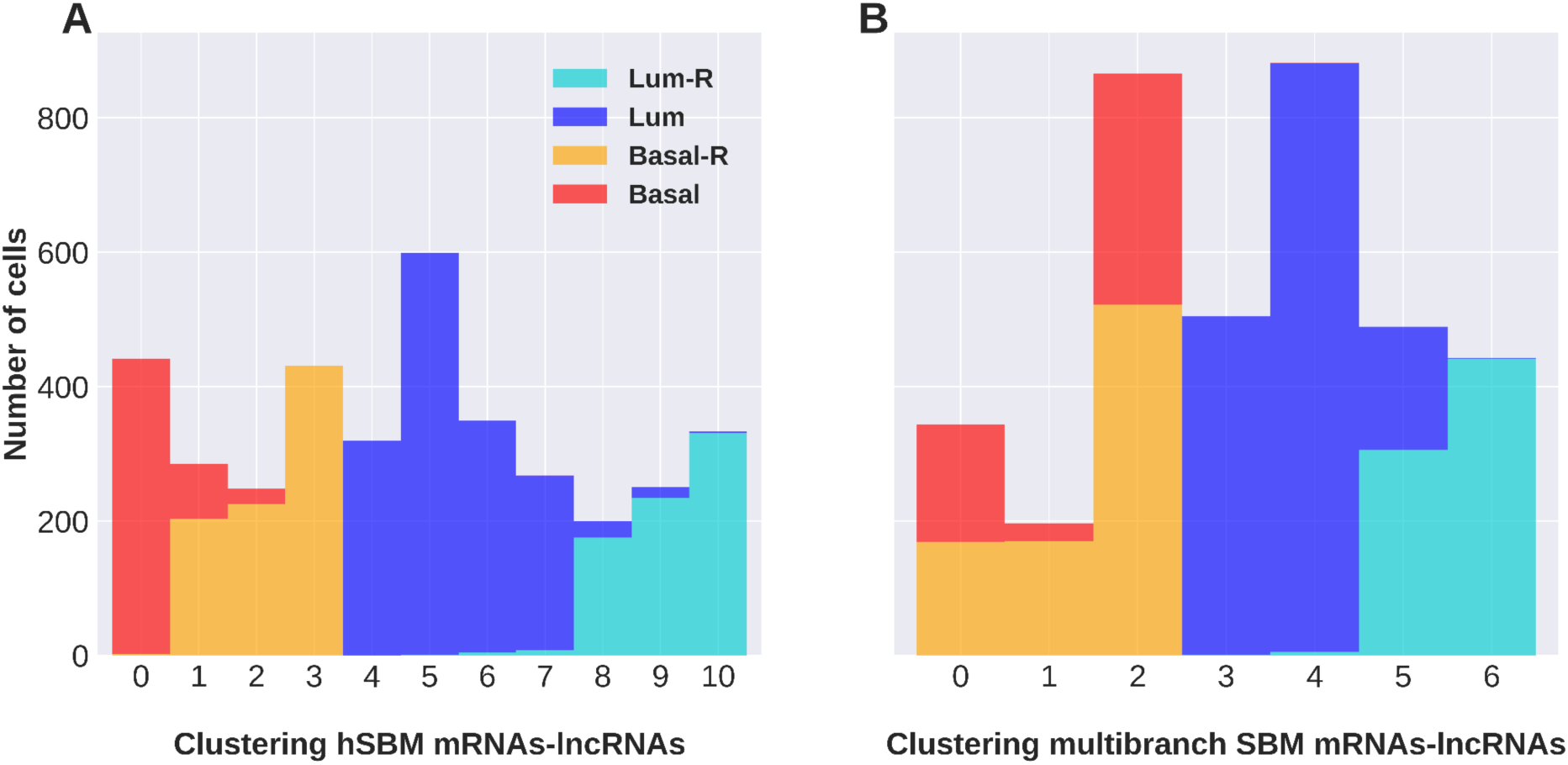
Clustering of cells obtained with hSBM applied to the concatenated mRNAs and lncRNAs expression matrices (A) and with the multibranch SBM applied to the mRNAs and lncRNAs matrices used as disjoint omics layers (B). Columns report the size of the identified clusters and the tumor of origin of their component cells.

The NMI/NMI* of level 1 obtained for the hSBM applied to the concatenated matrices is lower than the one obtained for mRNAs and lncRNAs analysed as disjoint omics layer (Table 1). The clustering obtained with the multibranch SBM algorithm, the NMI/NMI* score clearly improves compared to all previous experiments (Table 1), showing that mRNAs and lncRNAs matrices treated as disjoint omics layers provides more relevant information about tumor heterogeneity. In order to check whether the tri-partite approach leads to systematically better results than the bi-partite approach, we run multiple times each experiment (hSBM and multibranch SBM), measuring the NMI/NMI*. Each experiment was run 20 times and we excluded the runs with performances out of the 0.05 and 0.95 quantile in order remove outcomes with highly unlikely scores. Results of the multiple runs show that the multibranch SBM is statistically significantly better than the bi-partite approach (Figure 4).

**Figure 4.**
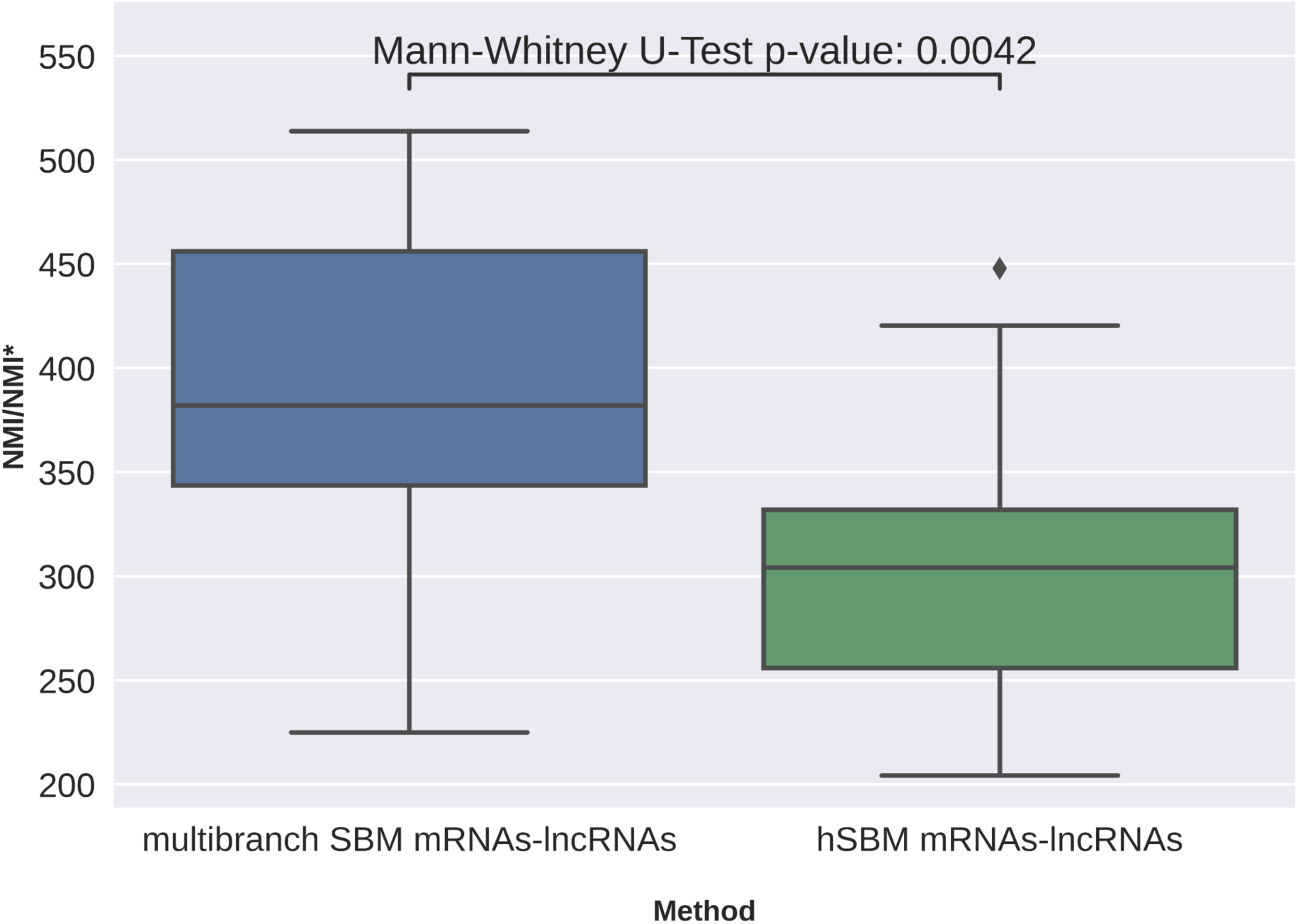
Boxplot showing the performances obtained with the multibranch SBM and with hSBM applied to the concatenated matrices.

### 3. Functional enrichment analysis of topics

We wanted to assess whether the gene content associated with the breast cancer cell clusters according to the multibranch SBM was enriched for specific biological functions. One of the pitfalls of enrichment analysis is the selection of input genes to be tested for enrichment [19]. The common practice in transcriptome analysis is to define a threshold to apply to the p-values to select differentially expressed genes. The cut-off is arbitrary and it has been shown that the effect of threshold choice impacts biological conclusions reached during enrichment analysis [20].

The hSBM and multibranch SBM algorithms provides in output topics that can be used as input lists for the enrichment analysis. Gene lists associated to topics are defined during the simultaneous optimal assignment of cells to clusters and genes to topics, without any dependency on an arbitrary threshold.

We tested through hypergeometric statistics the enrichment of topics for the gene functional annotations of the MsigDB Molecular Signature Database 7.5.1 (MsigDB) [21][22] and lncSEA database [23] for the mRNA-Topics and the lncRNA-Topics, respectively. We observed that the output of enrichment analysis is biased towards very long lists, therefore we decided to filter out uninformative gene sets before the test (see Methods). One of the two main innovation that we introduced is an unsupervised and reproducible procedure to select the most specific topics to each cluster, based on the probability matrix of association of a given topic to a given cell (see Methods). Results of topic enrichment analysis are provided in Figure 5.

**Figure 5.**
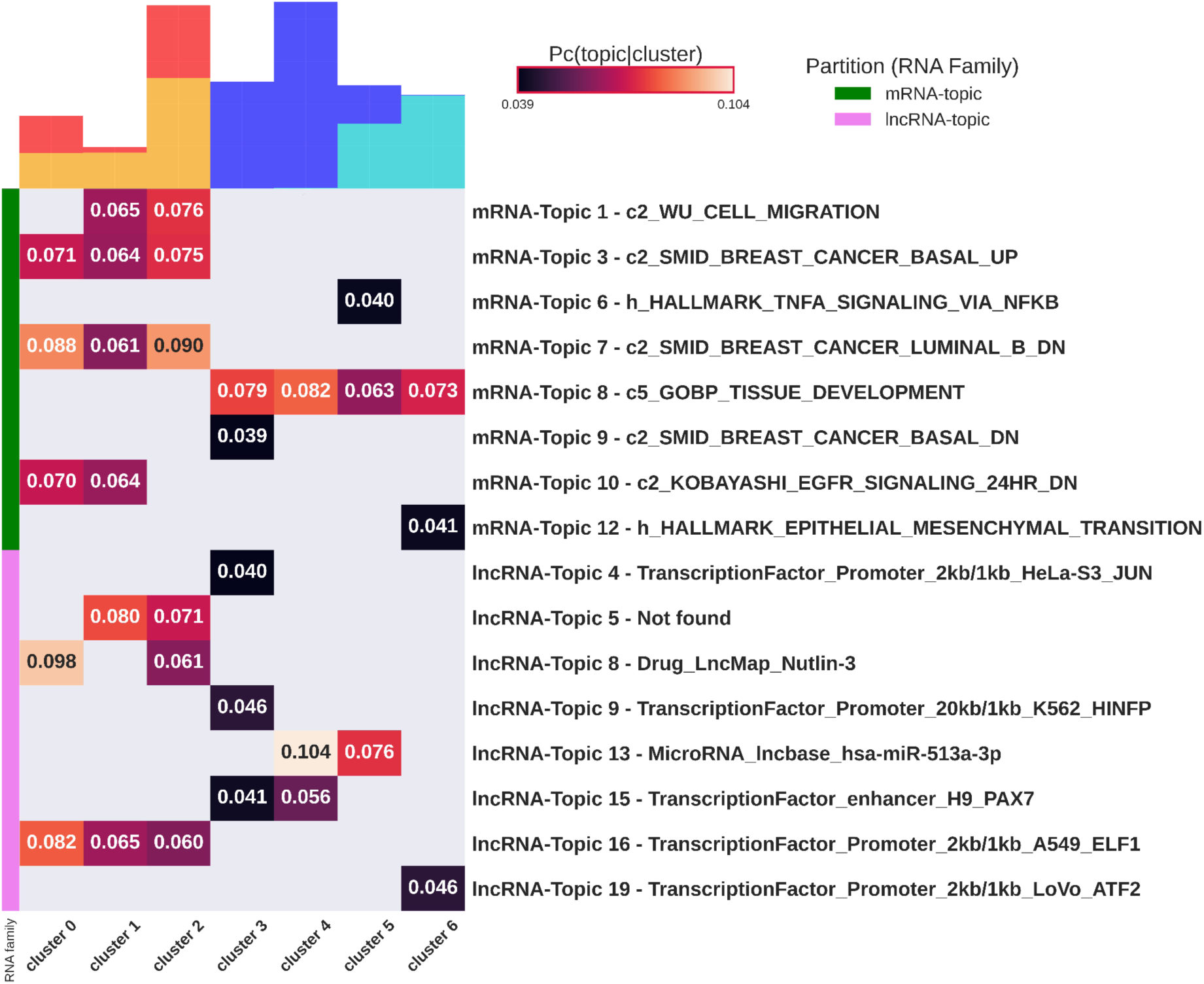
Functional enrichment analysis of topics. The heatmap reports the probability of association of topics to clusters (when statistically significant) and the most enriched functional gene. Green and pink lines highlight mRNA-Topics and lncRNA-Topics, respectively. On top of each column is shown the composition of the cluster in terms of tumor subtype color coded as in Figure 1 (red=“Basal”, yellow=“Basal-R”, dark blue=“Luminal” and light-blue=”Luminal-R”). All mRNA-Topics are associated to biological processes related to cancer and to gene signatures specific of breast cancer subtypes. The lncRNA-Topics are mainly associated to gene sets that represent interactors of transcription factors that play a role in cancer and in cell differentiation.

The results show a clear enrichment of all but one topic to the pathways involved in subtyping and progression of breast cancer and to sets of lncRNA that are interactors of transcription factors involved in cancer. Notably, we found topics specifically associated with luminal or basal cell clusters, with no topics associated with both luminal and basal types. As an example, mRNA-Topic 3 shows the keyword SMID_BREAST_CANCER_BASAL_UP [24] and is enriched only in basal-related clusters. The same happens for mRNA-Topic 7 which, being also composed of basal cells, shows a keyword related to luminal downregulation. All mRNA-Topics and lncRNA-Topics are associated to clusters of a specific tumor type and can be used in an unsupervised framework to identify the tumor type.

mRNA-Topic 1 is enriched in genes involved in cell migration and specifically associated to a subpopulation of cells from basal tumors, supporting a higher migratory capacity of basal cells compared to luminal ones [25]. Gene content in mRNA-Topic 6 is enriched in HALLMARK_TNFA_SIGNALING_VIA_NFKB and specifically associated to the cell cluster_5 containing a mixture of luminal resistant and sensitive cells. This result suggests the NF-kB regulation by TNFɑ as a transcriptomic hallmark common to a subset of resistant and sensitive luminal cells. Cell cluster_5 is also significantly associated with the lncRNA-Topic 13, which includes MALAT1 and NEAT1 as top ranked genes, two lncRNAs extensively studied in breast cancer progression [26] [27]. Another important result shows the association of cluster_6 composed only of therapy-resistant luminal cells, to mRNA-Topic 12 (HALLMARK_EPITHELIAL_MESENCHYMAL_TRANSITION) and lncRNA-Topic 19 (interactors of ATF2). The transcription factor ATF2 has been shown to induce therapy resistance to melanoma [28]. Finally, lncRNA-Topic 8, associated to two of the three basal clusters, consists of lncRNAs whose expression is correlated with the effect of Nutlin-3, a well known compound used in studies for cancer treatment [29]. In Table 3 we summarized some functional lncRNAs known in the literature that are associated to the identified topics.

### 4. Topic modeling versus clustering approach

In order to compare our method with a standard clustering approach, we decided to compare the topic modeling with clustering algorithm available in Seurat [30]. Seurat calculates cell clustering based on a single omic or with integrated omics with a weighted-nearest neighbour (WNN) approach [30]. We applied Seurat for clustering cells based on mRNA expression and lncRNA expression alone and then for integrated clustering (Seurat-WNN) with mRNAs and lncRNAs used as disjoint omics layers. We compared results obtained of Seurat clustering on individual omics with Seurat integrated clustering. To do this, we computed the adjusted mutual information (AMI) [31], a score based on mutual information as the NMI, but used when none of the two set of labels can be taken as ground truth. It varies between 0 and 1, being 1 the situation of maximal agreement between sets.

In terms of ability to reconstruct the correct tumor type, Seurat WNN achieves better performance as shown in table (Table 1). However, a deeper investigation showed that Seurat WNN does not fully exploit both omics, since the integrated clustering (Figure 6F) is very similar to the one obtained using mRNA expression alone (Figure 6B): both consist of eight clusters and each tumor type is separated into the same number of clusters. The agreement (AMI) between Seurat WNN and Seurat applied to mRNA alone is 0.855 (Table 2), confirming that introducing lncRNAs does not significantly change the results. The AMI between the Seurat clustering applied to lncRNA expression alone (Figure 6D) and Seurat WNN clustering (Figure 6F) is 0.569 (Table 2), which indicates very divergent results. We might impute this divergence to the cell specific modality weights calculated by Seurat WNN, that learns information content of each modality and determines its relative importance in downstream analyses [30]. Weights assigned to the mRNAs are significantly higher than those assigned to the lncRNAs (Figure 7), showing that mRNAs are predominantly used for the clustering.

**Figure 6.**
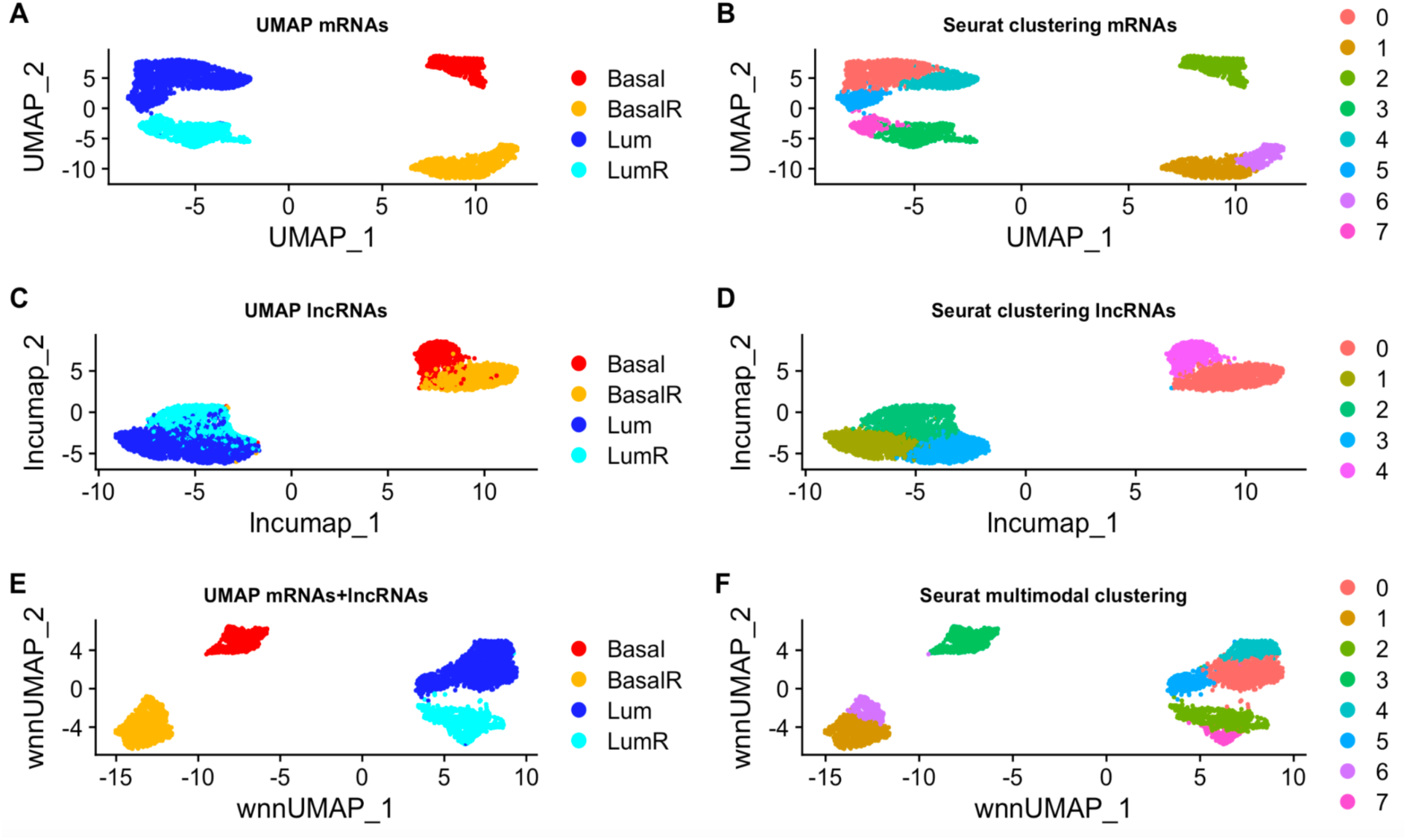
UMAPs representing the clustering obtained with Seurat using mRNAs expression alone (B), lncRNAs expression alone (D) and using both modalities (F). On the left is shown the tumor of origin of the cells in the UMAP embedding computed using mRNAs alone (A), lncRNAs alone (C) and the concatenated matrices (E).

**Figure 7.**
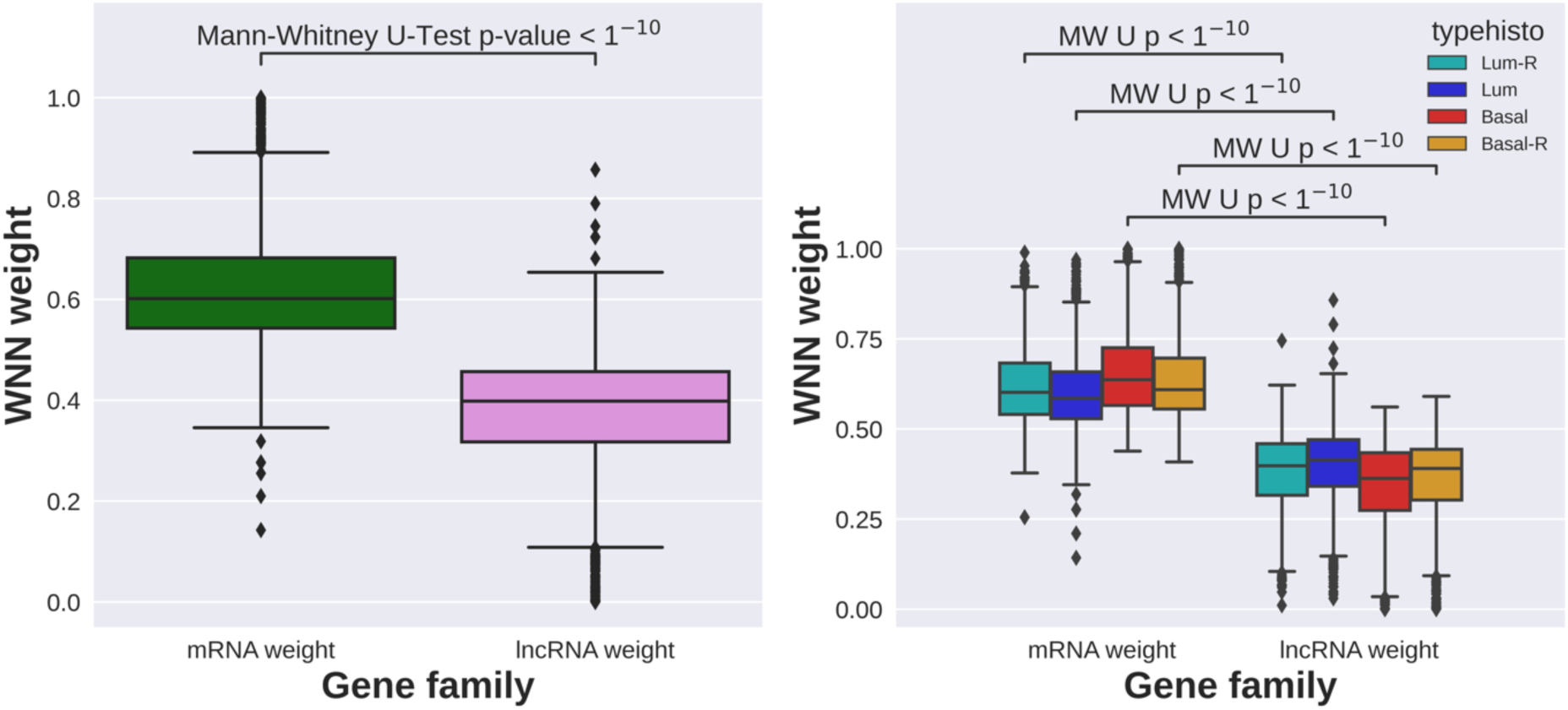
The violin plots show the weights assigned by Seurat WNN to the mRNAs and lncRNAs for all cells (A) and for cells grouped by tumor of origin (B). P-values are obtained with Mann-Whitney Utest.

**Table 2.** AMI scores computed to evaluate the concordance between the cell clusters derived from integrated clustering and single omics clustering.

| Experiment 1 | Experiment 2 | Adjusted Mutual Information |
| --- | --- | --- |
| Seurat-WNN<br>mRNAs-lncRNAs | Seurat-mRNA | 0.855 |
| Seurat-WNN<br>mRNAs-lncRNAs | Seurat-lncRNA | 0.569 |
| multibranch SBM<br>mRNAs-lncRNAs | hSBM-mRNAs | 0.492 |
| multibranch SBM<br>mRNAs-lncRNAs | hSBM-lncRNAs | 0.431 |

**Table 3.**
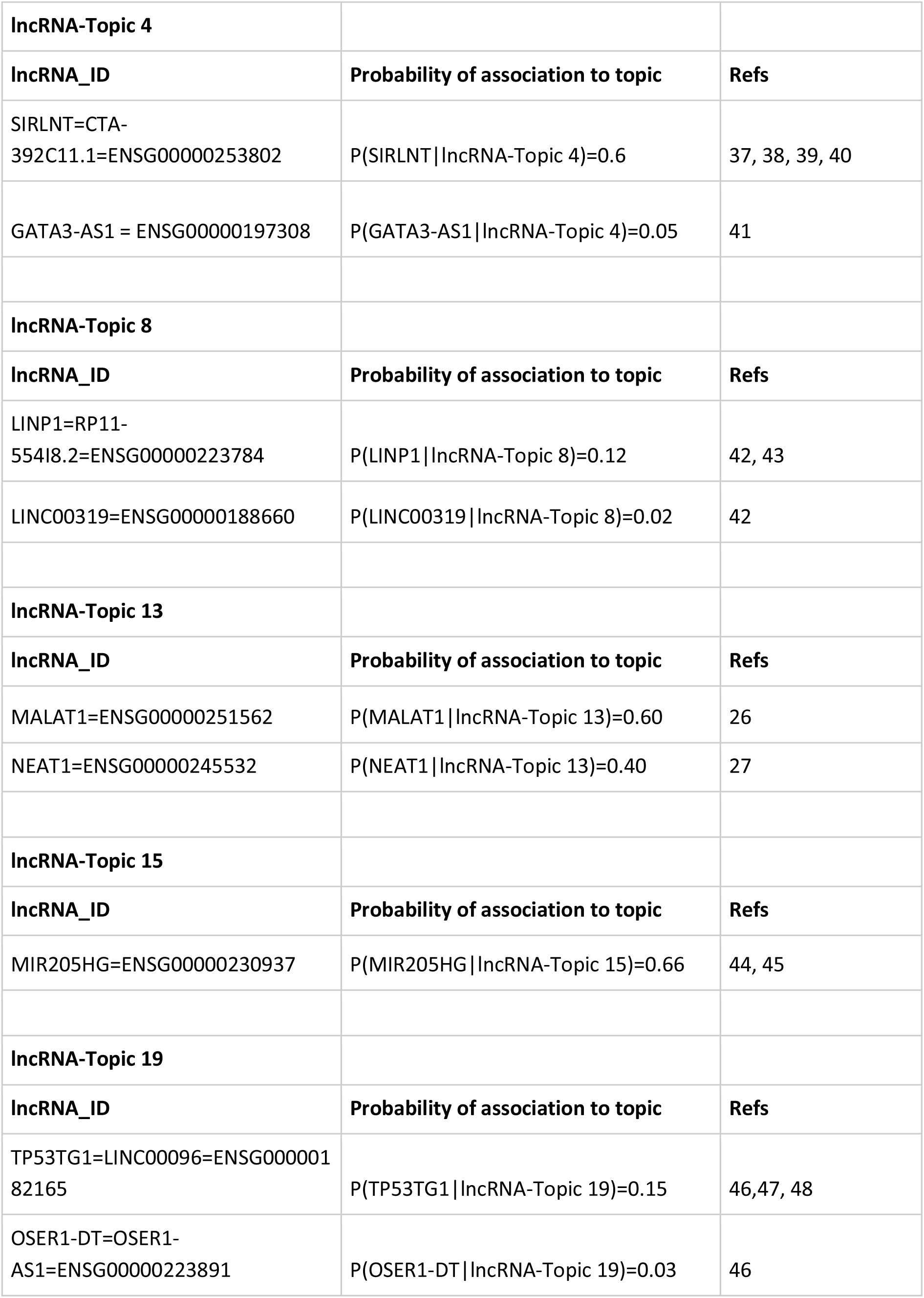
LncRNAs associated with lncRNA-topics with a high probability that have already been documented in the literature.

On the other hand, multibranch SBM assigns equal importance to each modality without introducing weights and possibly biases. The agreement between the integrated clustering and the clustering on the individual omics hSBM-mRNAs and hSBM-lncRNAs are, respectively, 0.492 and 0.493 (Table 2). This suggests that the proposed multi-branch SBM method, which does not rely on different weights associated with different omics, can be used as a complementary method to analyze single-cell multiomics data and extract information about cell subpopulations.

## DISCUSSION

The topic modeling procedure proposed here has proved to be a valid approach for integrative and unsupervised clustering of protein coding genes and lncRNAs expression at the single cell level. The multibranch SBM enables the classification of breast cancer cell populations, showing that mRNAs and lncRNAs datasets provide complementary information about the heterogeneity of the tumors under study, when they are treated as disjoint omics layers.

Combination of several single-cell omics layers represents a new frontier for single-cell genomics and requires appropriate computational methods. Recently, several experimental techniques are capable of assaying multiple modalities simultaneously from the same single cells, including CITE-seq [32], 10X Genomics ATAC + RNA, SHARE-seq [33], SNARE-seq [34] and many others. The multibranch SBM algorithm can be applied in different contexts of integrative multimodal analysis to learn the information content of each modality in each cell and to define cellular states based on a combination of information derived from multiple modalities.

Differently from state-of-art clustering methods used in single cell data analysis, such as Leiden algorithm and graph-based algorithms [35][36], multibranch SBM not only provides an integrative clustering of cells based on both mRNAs and lncRNAs expression, but also output the probability distribution of the association of these features to clusters, that is their relative importance in driving cell classification. Clusters of cells and sets of associated features (topics) are simultaneously identified with SBM algorithms. Topics can be used for functional enrichment analysis avoiding the arbitrary cutoff on the p-values to define input lists. Another great advantage of the SBM algorithm is that it automatically detects the optimal number of clusters and hierarchically clusters both the features and cells. This avoids researchers from fixing the number of expected clusters, which is difficult in real datasets. In addition, we show that clustering performed with multibranch SBM is not affected by batch effect, a common problem affecting the analysis of large-scale single-cell transcriptomic datasets (Supplementary Figure 4). The comparison with Seurat WNN has shown that we can better exploit the two omics layers whereas Seurat WNN relies more on the mRNA information. In our study of breast cancer PDX models, the highest hierarchical level separated the luminal and basal cells, while the second level showed a finer tumor heterogeneity.

Compared to standard state-of-art clustering analysis, our method provides simultaneous optimal partitioning of genes into topics and cells into clusters, allowing for interpretable clustering results. We proposed a new approach to investigate the heterogeneity of single-cells, based on a reproducible and unsupervised method to associate to each cluster the set of topics that identifies the behavior of the cells within a cluster, that remove the arbitrariness in the interpretation of the enrichment analysis. Identified topics associated to clusters are enriched for genes and pathways clearly involved in breast cancer subtyping, such as cell adhesion and migration and in sets of lncRNAs that are interactors of transcription factors related to cancer.

Overall, we identify some lncRNAs with high probability of association to topics that are well known in breast cancer literature and some others very interesting candidates for functional validation. In the lncRNA-Topic 4, which includes a set of 19 lncRNAs, SIRLNT (SIRT1 regulating lncRNA tumor promoter) has the top high probability of association to the topic (P(SIRLNT|lncRNA-Topic 4)=0.6). This lncRNA, also called lncRNA-PRLB (progression-associated lncRNA in breast cancer), is upregulated in human breast cancer tissues and breast cancer cell lines [37]. Moreover, it has been described as a modulator of SIRT1 gene both at the mRNA level and protein level. SIRT1 is a member of the sirtuin family of proteins extensively studied in multiple cancers [38], including breast cancer [38][40].

Also strongly associated with lncRNA-Topic 4 we found GATA3-AS1, a lncRNA recently proposed as a predictive biomarker of nonresponsive breast cancer patients to neoadjuvant chemotherapy [41]. GATA3-AS1 was described as a functional lncRNA and positive transcriptional regulator of *GATA3* in TH2 lineage of lymphocytes [41].

The lncRNA-Topic 8 is strongly associated with clusters 0 and 2, both of which are composed of triple negative sensitive and resistant cells. LncRNA-Topic 8 presents among its top features some lncRNAs extensively studied in breast cancer, and particularly in triple negative subtype. The top ranked lncRNA in lncRNA-Topic 8 is LINP1, a regulator of DNA repair in triple negative breast cancer [42] whose knockdown increases the sensitivity of the breast cancer cells to radiotherapy [43]. A second relevant lncRNA belonging to lncRNA-Topic 8 is LINC00319, which is been shown to promote cancer stem cell-like properties [42].

LncRNA-Topic 13 is characterised by two lncRNAs associated with it with a very high probability that are well known in the literature, namely MALAT1 (Metastasis Associated Lung Adenocarcinoma Transcript 1) and NEAT1. MALAT1 is one of the most studied non-coding RNA in cancer, generally overexpressed in tumor progression and metastasis [26]. NEAT1 is also abnormally expressed in many cancers and associated with therapy resistance and poor clinical outcome [27]. In our analysis, the lncRNA-Topic 13 is significantly associated with cluster 5 which contains a mixed subpopulation of sensitive and therapy-resistant luminal cells, suggesting that a subset of sensitive cells share this lncRNA signature with resistant cells.

The host gene for miR-205 (MIR205HG) is the most strongly associated with lncRNA-Topic 15, expressed in luminal sensitive cell cluster 1 and cluster 2. MIR205HG has independent functions as a lncRNA, regulating growth hormone and prolactin production in the pituitary gland [44]. Remarkably, MIR205HG was previously indicated as a predictor of anti-cancer drug sensitivity [44][45].

The LncRNA-Topic 19 is specifically expressed in cluster 6 of luminal drug-resistant cells. The top ranked lncRNAs associated with lncRNA-Topic 19 are LINC00493, also known as SMIM26, and TP53TG1. LINC00493 and TP53TG1 are enriched and co-released in extracellular vesicles from colorectal cancer cells [46]. Previous studies have highlighted the ambivalent oncogenic or tumor suppressor activity of TP53TG1 in luminal breast cancer [47][48].

## METHODS

### scRNA-seq dataset pre-processing

Single cell expression count matrices were obtained from the NCBI GEO repository (GSE117309). This dataset includes scRNA-seq profiles from 3723 cells of patient-derived xenograft breast cancer models. Cells were collected from four cancer models: a luminal untreated drug-sensitive tumor, a luminal breast tumor with acquired resistance to Tamoxifen, a triple negative tumor initially responsive to Capecitabine and a triple negative tumor with acquired resistance to Capecitabine.

Protein-coding mRNAs and lncRNAs raw expression matrices were obtained based on Ensemble grch37 gene annotations [49], after removing mitochondrial and ribosomal genes. Data were analyzed with Scanpy version 1.8.2 [50]. Library size normalization was applied and highly variable mRNAs and lncRNAs were selected according to the Scanpy procedure, selecting 1959 and 1136 features respectively. We utilized two distinct thresholds for the parameter min_disp, considering the significant difference in the intrinsic coefficient of variation between the expression levels of the two gene families, as observed in our dataset (Supp. Fig 1).

### hSBM algorithm

We used the original hSBM clustering algorithm available at https://github.com/martinger-lach/hSBM_Topicmodel/ for topic modelling. The algorithm maximises the probability that the model *M* describe the data *A*. Formally it maximises the quantity *P*(*A*) = *P*(*M*)*P*(*M*). In our context, *A* is the gene expression matrix and the model *M* is composed of the levels of the hierarchy, each with its own partitions of genes and cells. hSBM minimises the description length of the model, defined as ∑ = −*lnlnP*(*M* ∨ *A*). The most important hyperparameter in the hSBM implementation of [6] is the number of initializations, i.e. the number of different starting points from which the algorithm begins to estimate the posterior probability *P*(*A*). We decided to run hSBM with seven initializations for each experiment because we found that seven initializations best balanced the time cost and the ability to reach the minimum description length (see Supp. Fig. 2A). Multibranch SBM is the multipartite extension of hBSM: it works with a multipartite graph and finds partitions on each side of the graph [4]. We also ran Multibranch SBM with seven initializations since it also shows an elbow at seven initialisations (Supp. Fig 2B).

### Multibranch SBM algorithm

The analytical framework for adding multiple layers of information to a hSBM inference problem was recently introduced for text analysis [51]. In the context of scRNA-seq data analysis, the algorithm was adapted to handle a tri-partite network where nodes represent mRNAs, lncRNAs and cells. mRNAs and lncRNAs were connected to cells according to their expression level, therefore there is no edge between the mRNAs and lncRNAs nodes. The statistical inference procedure, as well as the definition of topics and probability distributions, leading to the tri-partite network clustering is an extension of the hSBM algorithm. The n-partite SBM, introduced in [4] extends the hSBM algorithm described in [6]. Its implementation is available on GitHub at https://github.com/BioPhys-Turin/nsbm.

The inference procedure is performed using the functions implemented in the graph-tool library [52]. This algorithm infers the blocks structure on a given network, using a Markov Chain Monte Carlo (MCMC) that moves nodes (i.e. genes, lncRNAs, cells) between blocks in order to minimize the so-called Description Length (DL) ∑ [53]; this represents, in nat units, the number of bits that the model needs to describe the data. It can be written as the logarithm of the product of the likelihood times the prior ∑ = *ln*,*P*(*A* ∨ *M*)*P*(*M*)(*A* is the matrix of the data, *M* represents all the model parameters) [54].

This description length ∑ can be optimised [55] by a MCMC whose moves are accepted with the following probability:

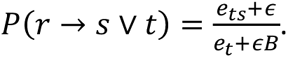

The probability of moving a node from block (cluster or topic) r to a blocks, given that a random neighbour of the considered node belongs to block t is proportional to the ratio between the number of edges that connects group t and blocks *e*_*ts*_ and the number of edges that go to block t *e_t_* plus a term ∈*B* that is necessary to have a uniform probability if there are no connections between the blocks considered.

hSBM and the multibranch SBM impose that nodes of a given type (mRNAs, lncRNAs, cells) must not be mixed, so the blocks can contain only homogeneous nodes. This allows us to define clusters (blocks of cells), mRNA-Topics (blocks of mRNAs) and lncRNA-Topics (blocks of lncRNAs).

Once the model is run, moving all the nodes multiple times and accepting or refusing the moves with the aforementioned probability, it is possible to extract the P(mRNA | mRNA-Topic) and P(mRNATopic | cell) as well as P(lncRNA | lncRNA-Topic) and P(lncRNA-Topic | cell)

The P(mRNA | topic) is the ratio of the number of edges connected to the mRNA and the number of edges connected to the topic (the total number of edges connected to any gene in the topic). P(topic | cell) is the ratio of the total number of edges connected to the cell and number of edges connecting the cell to the topic. The same definitions apply to P(lncRNA | lncRNA-Topic) and P(lncRNA-Topic | cell).

Finally, let us highlight that looking at the probability of accepting a move *P*(*r* → *s* ∨ *t*) it is clear the advantage of using multibranch SBM on multi-omics problems: when moving a cell from block r to block s the algorithm samples a neighbour of the cell and it labels its group as t. At this point the model only considers the links between the cell and the nodes in group *t*, so the normalisations of other branches are irrelevant. In other words, at each move cells are clustered looking at a single omics and this allows to naturally perform inference on multi-omics datasets in which each omic has a different normalization.

### Normalized Mutual Information score

To evaluate clustering performance, we computed the Normalized Mutual Information (NMI) as defined in [56] between the partition found by the algorithm and the tumor of origin of the cells. NMI is a widely used measure to compare clustering methods [57]. However, the need for adjustment for the so-called selection bias problem is needed, that is NMI steadily increases with the number of clusters. To keep this effect into account, we adjusted the empirical NMI with the NMI* obtained with a null model that preserves the number of clusters and their sizes but reshuffles the labels of samples. Thus, NMI/NMI* represents how much the empirical score is higher than the score obtained with random populated clusters of the same size and number.

### Topic to cluster assignment

We developed a procedure to assign to each cluster its more suitable topics, exploiting the *P*(*topic* ∨ *cell*).

First, we computed the *P*_c_(*topic* ∨ *cell*) because it has been shown that is more informative than the *P*(*topic* ∨ *cell*) (11). This is defined as

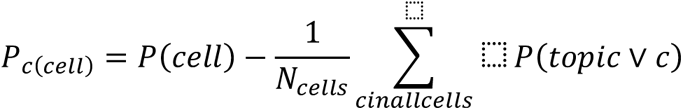

Then, we calculated the **P*_c_*(*topic* ∨ *cluster*) as mean of *P*_c_ (*topic* ∨ *cell*) over the cells inside the cluster

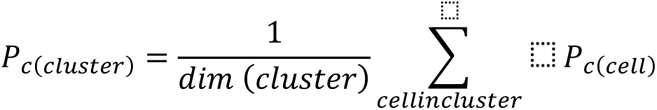

Finally, we assigned to each cluster all the topics that satisfied the condition

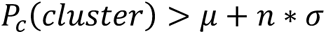

where µ and σ are the mean and the standard deviation of the **P*_c_*(*topic* ∨ *cluster*) across all the topics for the cluster under examination.

The core of the procedure concerns the choose of *n*: to make it unsupervised and reproducible we started from *n* = 3, then we lowered *n* by step of 0.05 until two conditions were fulfilled; first we required that at least one topic is assigned to each cluster, second that there were no clusters with the same sets of topics, but subsets were allowed. It’s worth noting that following this method it’s possible to assign to any kind of partitions the most important topics for each subset: for example, one could group cells by subtype and, computing the *P*_’_(*topic* ∨ *subtype*), could identify the most important topics for each tumor subtype.

### Functional enrichment of topics

The gene content of topics was tested through hypergeometric statistics for functional enrichment using annotations from the MsigDB Molecular Signature Database version 7.5.1 (MsigDB) [21][22] and the lncSEA database [23]. We tested all topics assigned to the clusters, excluding those not associated with any of the cell partitions. After correcting hypergeometric test p-values for multiple testing using Benjamini-Hochberg [58], we associated each topic to the most enriched functional gene set. When a gene set is associated with multiple topics, the one with the lowest false discovery rate was retained.

We observed a strong bias in favor of large size gene sets as outcome of the hypergeometric test when using the lncSEA database: most of the genes in the topics corresponded to the collection called “Accessible Chromatin”, more precisely, to gene sets consisting of more than 12000 genes.

To investigate the impact of these large size gene sets on our results, we studied the structure of MsigDB and lncSEA databases. MSigDB contains 32880 gene sets divided in 9 collections including total of 40755 genes. Collections correspond to set of genes constructed by the same method or exploring similar biological information. For example, the collection “Hallmarks” (H) of MsigDB consists of “coherently expressed signatures derived by aggregating multiple gene sets to represent well-defined biological states or processes” [21]. LncSEA contains 41365 gene sets divided into 18 collections and 58478 genes. We computed the ratio between each set size and the total database size (the total numer of genes in the database). The distributions of the ratios calculated for the two databases have a similar shape but the maximum value of the ratio for MsigDB is 0.05 while for lncSEA it is 0.25 (Suppl Fig. 3A, B). This suggests that the lncSEA database comprises numerous gene sets with a large number of elements, ranging from 10000 to 14000, that may have questionable biological significance. To filter out extremely long gene sets, we calculated the ratio between the size of each gene set and the total number of genes within the collection which the gene set belongs to. All gene sets whose ratio was greater than 0.15 were filtered out.

Nevertheless, this filtration method fails to tackle the problem of redundant gene content present within sets. To eliminate redundant sets, one possible solution is to calculate the Jaccard distance between each pair of gene sets and remove those that are more similar than a specific threshold. For a collection with n gene sets, this requires computing n(n+1) operations, as each set’s intersection and union must be calculated with all others, resulting in 2*n(n+1)/2 operations. However, collections such as C11 and C13 in the lncSEA database contain around 14,000-18,000 genes, which makes this operation incredibly time-consuming and computationally demanding. When dealing with a total of approximately 70,000 gene sets between the two databases, this method becomes almost impractical. Nevertheless, we can demonstrate that, for a subset of classes, the Jaccard distance and gene set length are linked by a decreasing monotonic relationship. In other words, the longer a list, the more similar it is to other lists in the same class. Consequently, implementing the intersection-with-universe filter is a quick and accurate approximation of the Jaccard filter (Suppl. Fig 3C, D, E, F).

## AVAILABILITY

Scripts to reproduce all the analyses and figures presented in this paper are provided in the GitHub repository https://github.com/sysbio-curie/topic_modeling_lncRNAs.

## SUPPLEMENTARY DATA

Supplementary Data are available online.

## ACKNOWLEDGEMENTS

We would like to acknowledge the Competence Centre for Scientific Computing C3S which provided us the access to the computing cluster OCCAM [59]. This work was supported by the European Commission’s Horizon 2020 Program, H2020-SC1-DTH-2018-1, “iPC individualizedPaediatricCure” (ref. 826121).

## Author Contributions statement

L.M. and M.C.: conceptualization, methodology, G.M. and F.V.: methodology, data curation, data analyses, formal analysis. L.M. G.M and F.V. Writing-original draft and editing. All authors reviewed the manuscript.

## Competing Interests statement

The authors declare no competing interests.

---

**Supplementary Figure 1.**
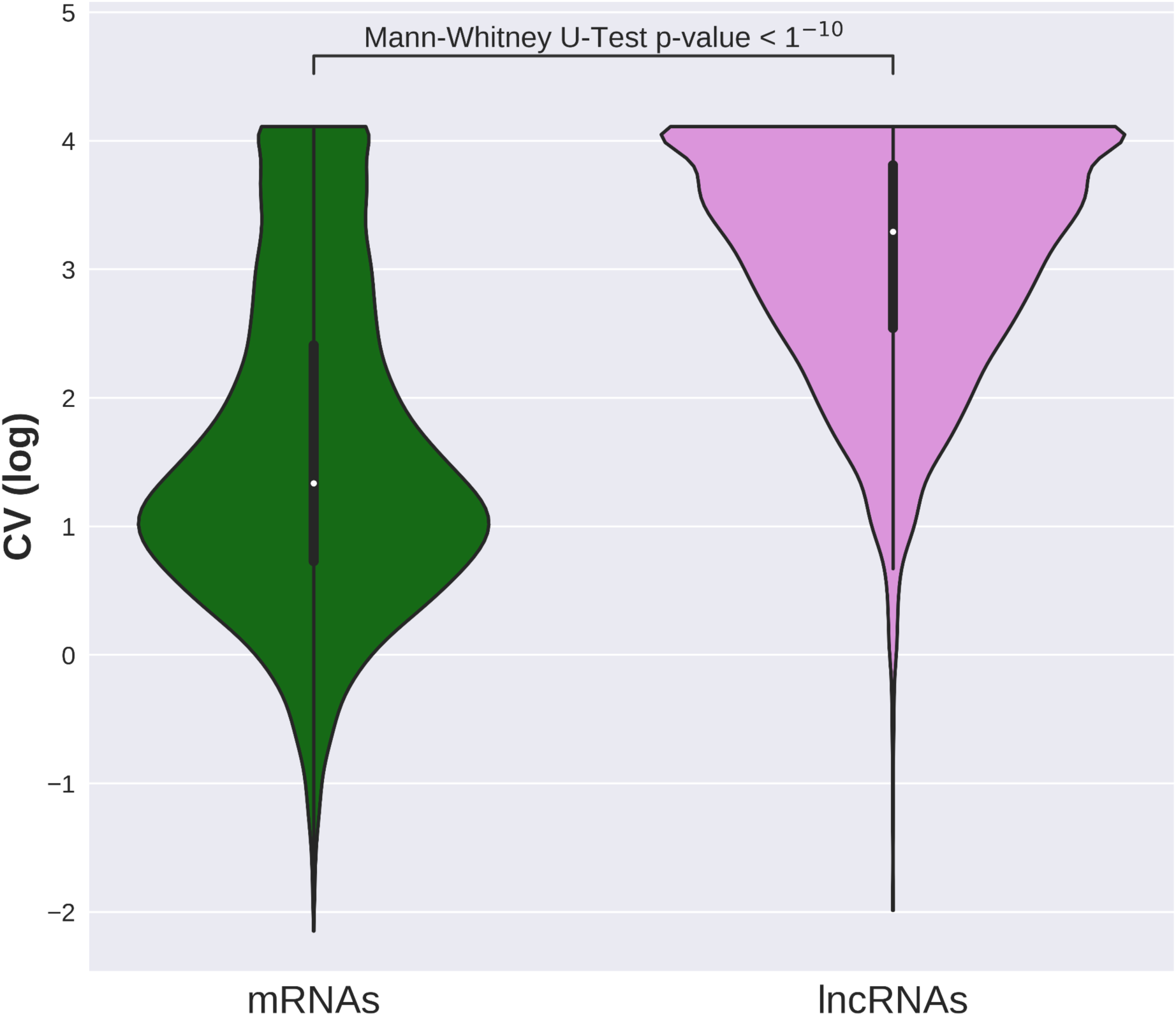
Violin plots showing the distribution of the coefficient of variation of the mRNAs (green) and lncRNAs (pink). The p-value indicates the outcome of the Mann-Whitney U-Test.

**Supplementary Figure 2.**
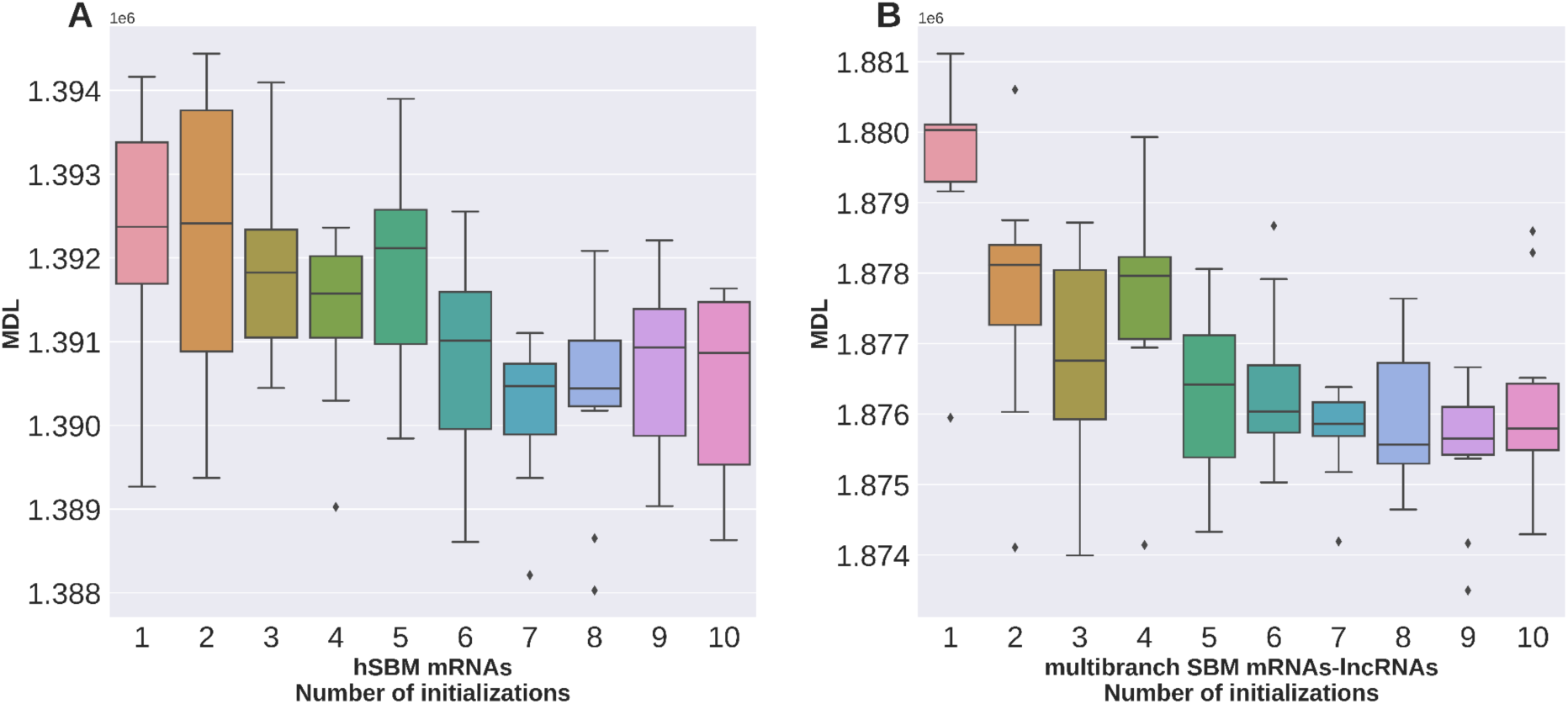
Boxplot of the minimum description length (MDL) reached varying the numbers of initializations for the experiment hSBM-mRNA (A) and the multibranch experiment (B). Both hSBM and multibranch SBM show an elbow at seven initializations.

**Supplementary Figure 3.**
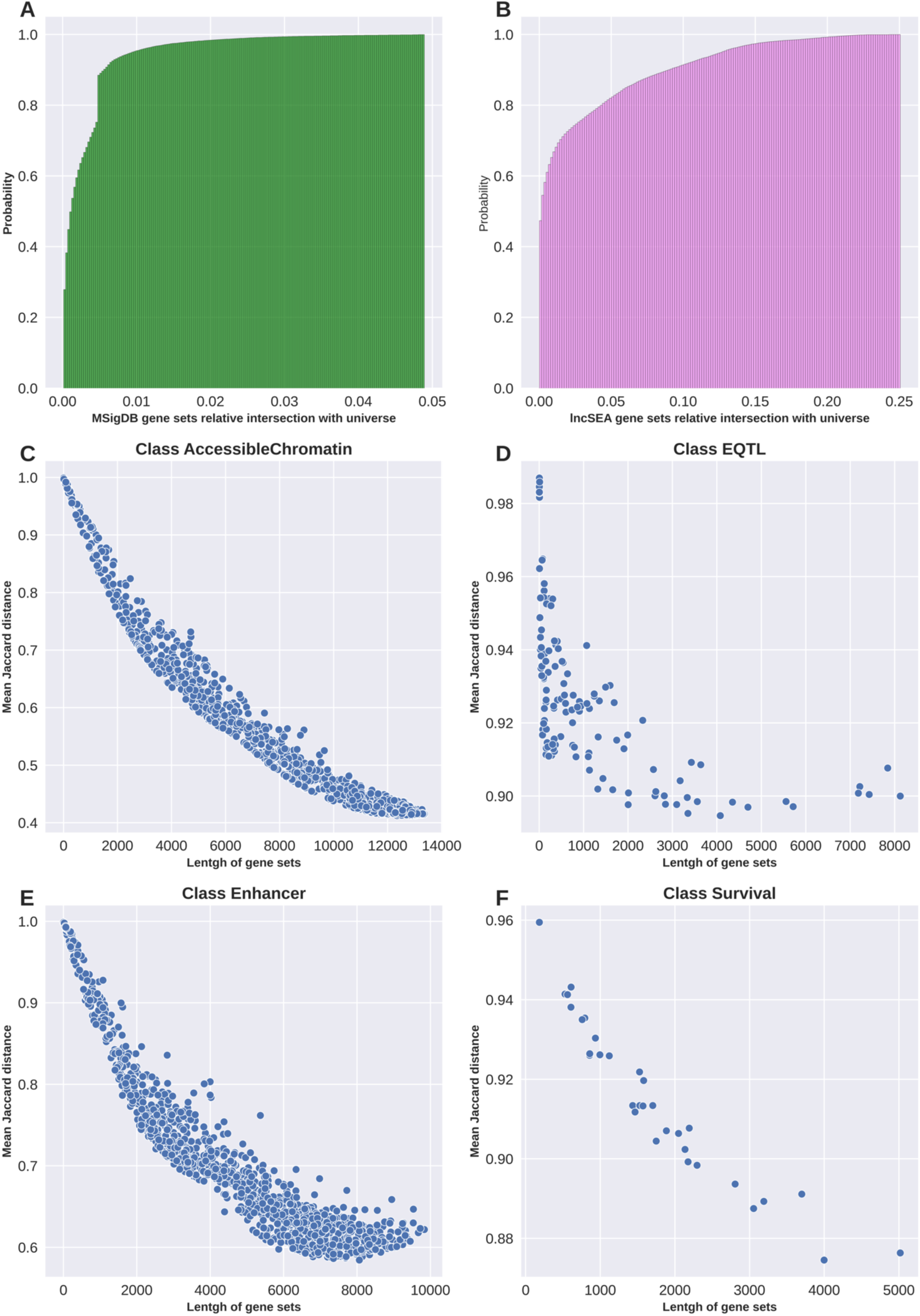
Distribution of the length of the gene sets in the MSigDB database for the mRNAs (A) and lncSEA database for lncRNAs (B). The latter consists of very longer gene sets in comparison to the former. Four examples of the relationship between the length of the gene sets and the mean similarity, measured with the Jaccard distance, among the sets (C, D, E, F). The decreasing monotonic relationship allows for the using of the filter based on the length instead of the Jaccard distance since the latter is highly computationally demanding.

**Supplementary Figure 4.**
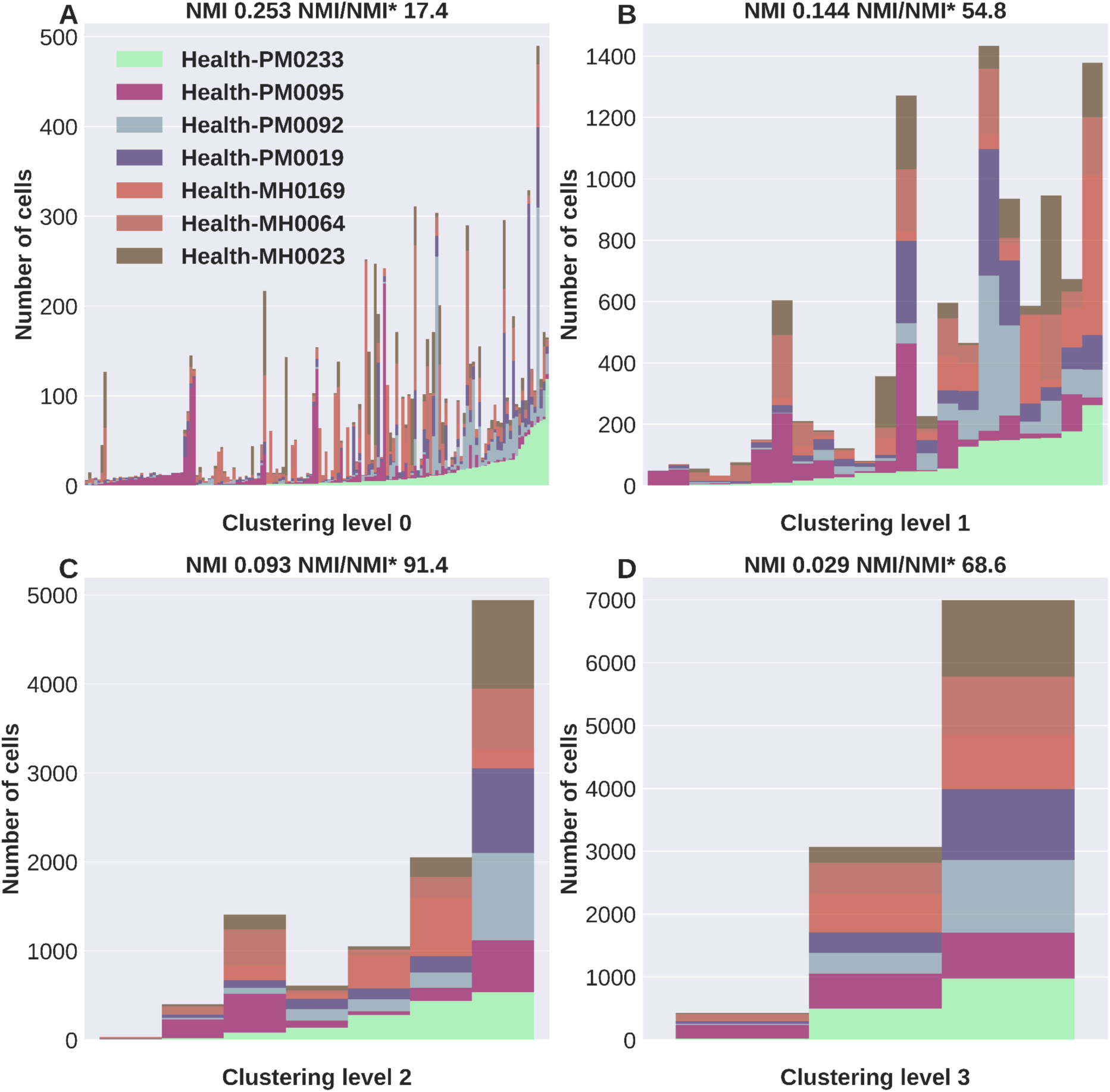
Hierarchical clustering of cells from seven healthy donors of breast tissue. Each panel shows the clusters belonging to one level of the hierarchy. We applied the multibranch algorithm creating the three usual partitions: cells, mRNAs and lncRNA. The low values of NMI and NMI/NMI* and the homogeneous composition of each cluster prove that the algorithm cannot recognize the donor.

## Notes

### Competing Interest Statement

The authors have declared no competing interest.

### Summary of Updates

Discard a comment accidentally included in the main text.

