## Supplementary figures and images for "Identification of interpretable clusters and associated signatures in breast cancer single cell data: a topic modeling approach"

### Supplementary Figure 1

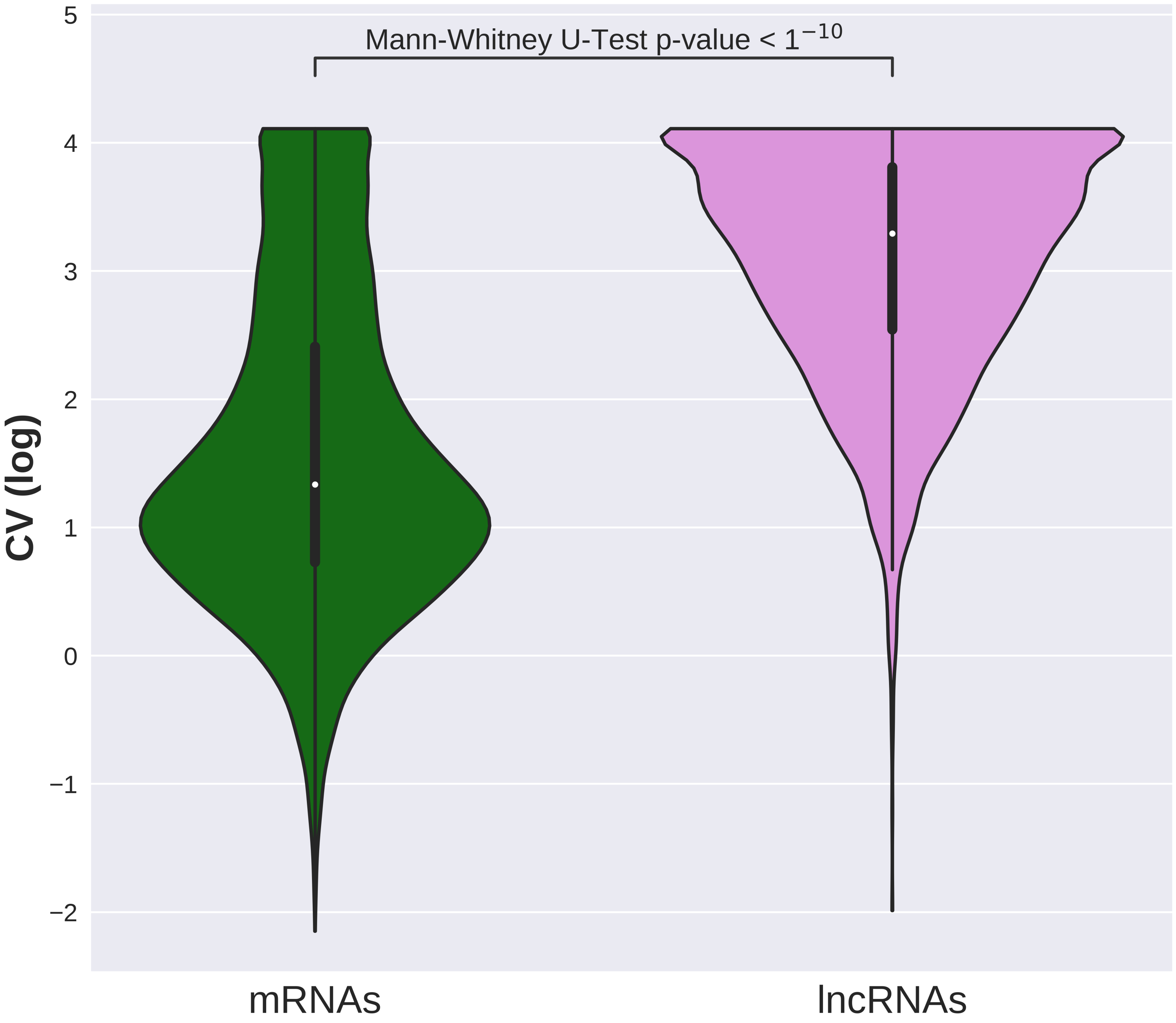

### Supplementary Figure 2

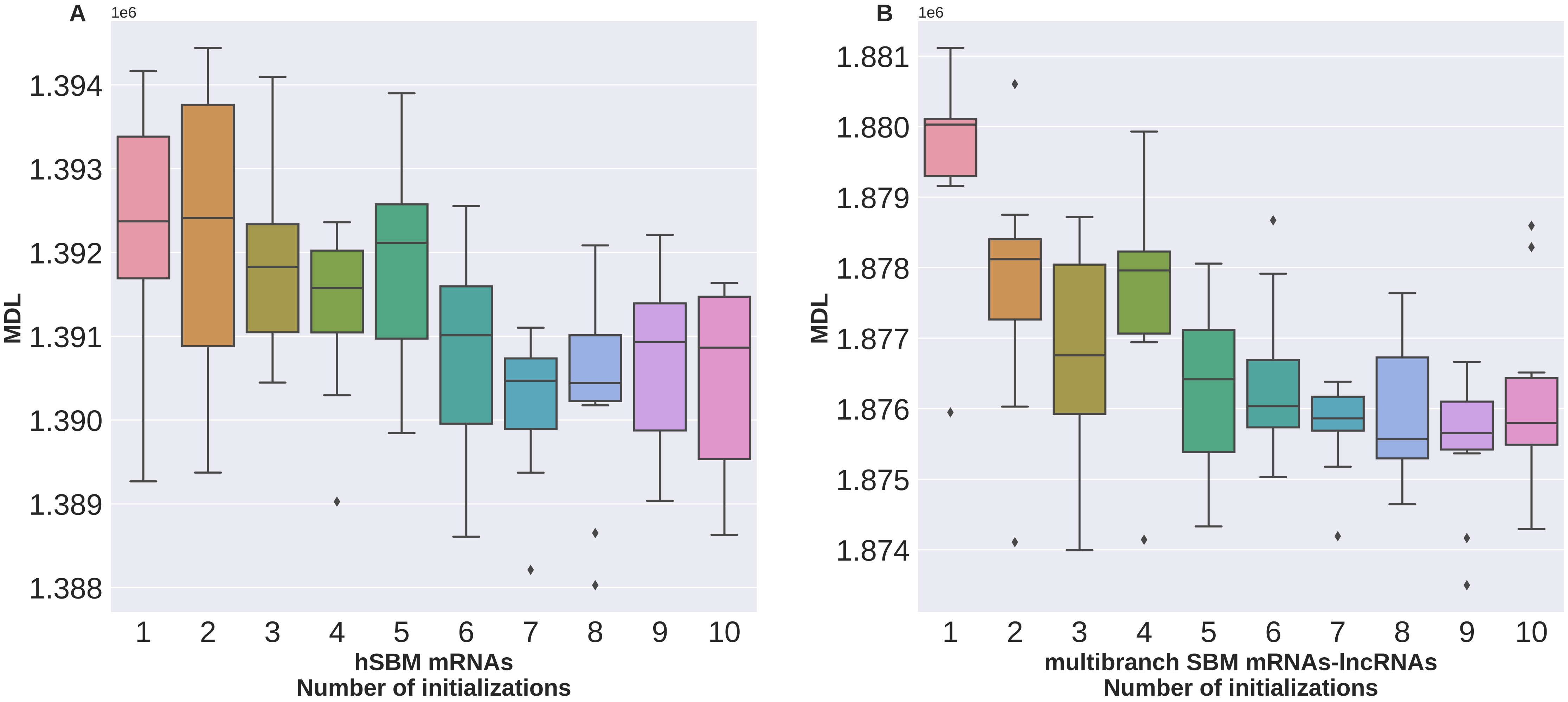

### Supplementary Figure 3

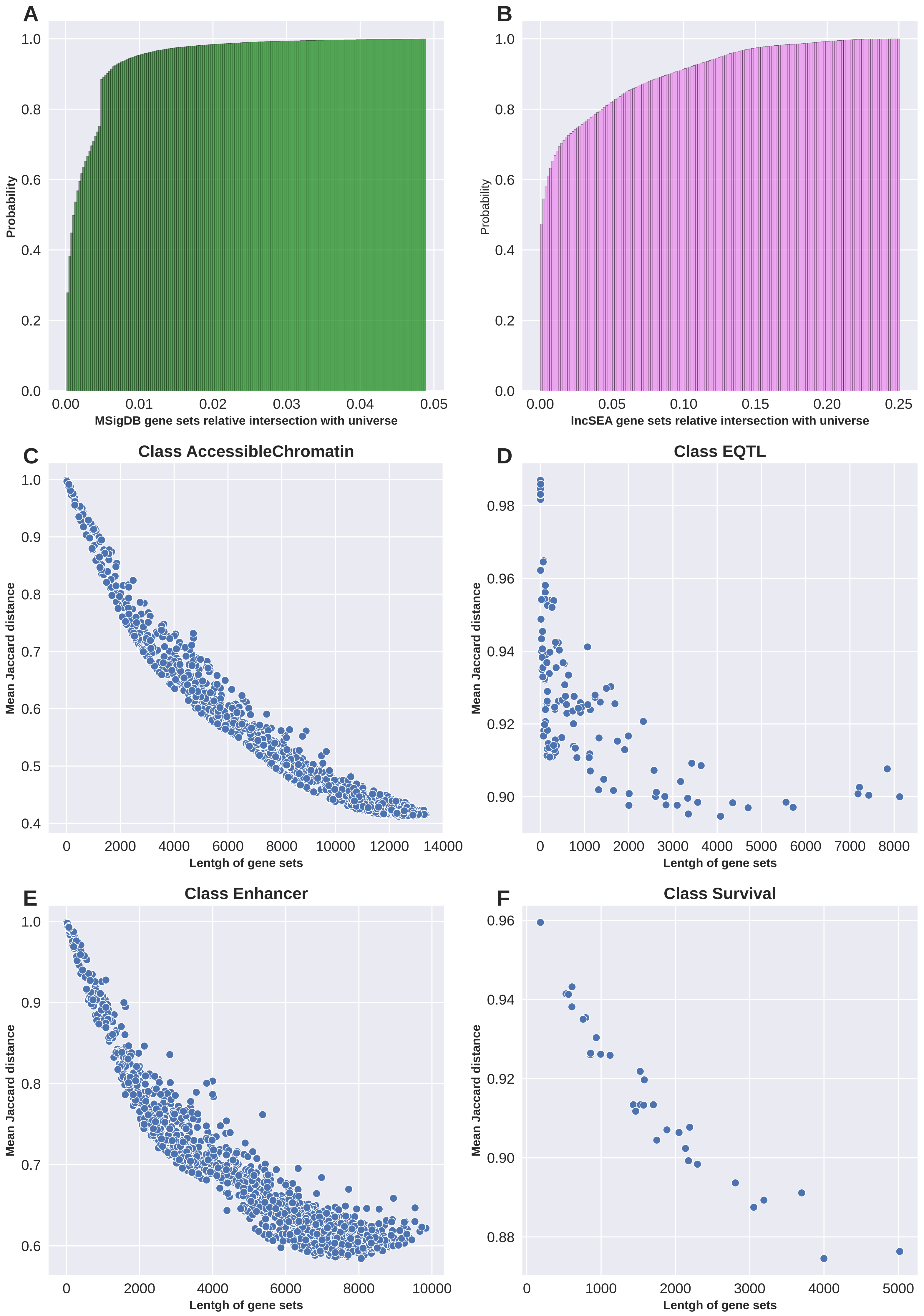

### Supplementary Figure 4

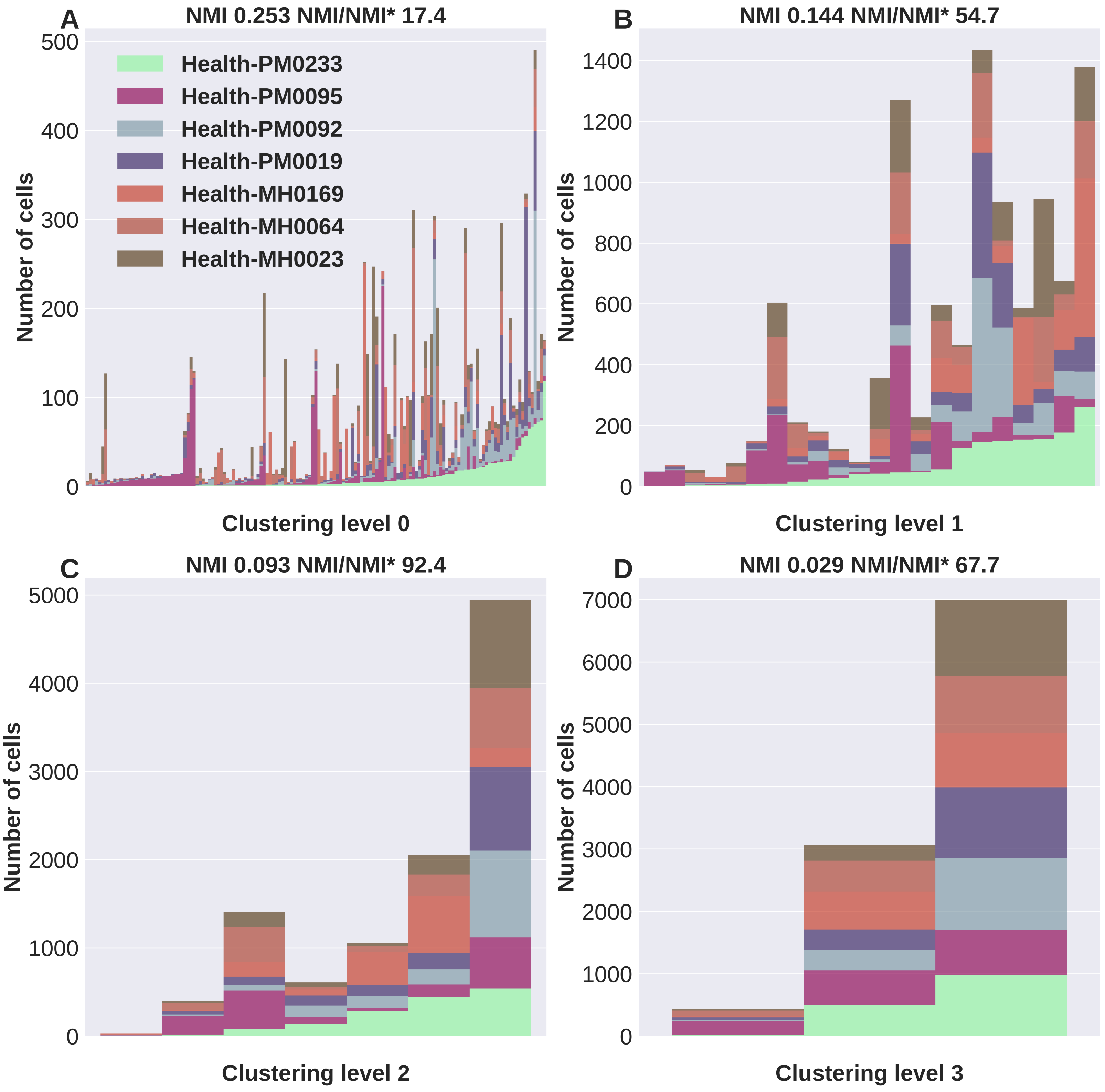
